# DNAi: an open-source AI tool for unbiased DNA fiber analysis

**DOI:** 10.1101/2025.09.30.679603

**Authors:** Clément Playout, Yosra Mehrjoo, Renaud Duval, Marie Carole Boucher, Santiago Costantino, Hugo Wurtele

## Abstract

DNA fiber assays are powerful tools for investigating replication dynamics at the single-molecule level. However, their application and widespread adoption has been hampered by the labor-intensive and tedious nature of manual analysis of large numbers of images. Quantification of labeled DNA fibers typically depends on subjective examination, selection, and annotation of individual fibers from fluorescence microscopy images reducing inter-user consistency, reproducibility, and experimental throughput. To address these issues, we developed DNAi, a computer vision tool based on deep learning allowing automated detection and quantification of labeled DNA fiber length. DNAi was trained on a large and diverse dataset of manually annotated images of DNA fibers and matches human performance and accuracy in segmentation and length measurement across a wide range of experimental conditions. The open-source tool includes a user-friendly interface, which permits visual validation and manual selection of segmented fibers. Overall, DNAi enables robust, rapid, and reproducible DNA fiber analysis, and is freely available.

**Graphical abstract:** 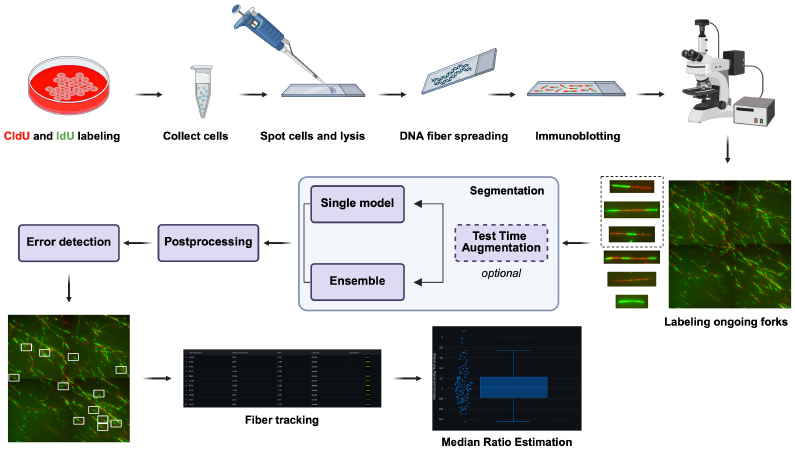

## Introduction

During the S phase of the cell cycle, cells undertake the critical task of faithfully and completely replicating their genetic material. DNA replication is initiated at chromosomal sites called “origins”, from which two replication forks originate and move in opposite directions along the DNA molecule (reviewed in [3, 15, 6]). Impediments to DNA replication fork progression can be caused by a plethora of environmental or endogenous genotoxic agents, resulting in a state of replicative stress [19, 13, 34, 30, 14]. Interestingly, several developmental syndromes, e.g., primordial dwarfism, have been shown to be caused by mutation in DNA replication factor-encoding genes or in mediators of the replicative stress response [17, 5, 23, 16, 22, 25, 18]. Moreover, replicative stress is a well-known driver of cancer-associated mutagenesis and response to genotoxic chemotherapy drugs [10, 9, 13]. Accordingly, characterizing DNA replication dynamics has become the focus of intense research interest.

Experimental studies of DNA replication have relied extensively on labeling nascent strands with halogenated deoxyuridine analogs, including chloro-(CldU), bromo-(BrdU), and iodo-deoxyuridine (IdU). Cells are incubated with media containing such exogenous nucleoside analogs, which are rapidly incorporated in replicating DNA. The dynamics of incorporation of these analogs can then be analyzed using immunofluorescence (IF) strategies combined with flow cytometry or microscopy. Among those methods, DNA fiber spreading and combing [4, 27, 8], rely on the stretching of individual DNA molecules on glass slides or coverslips, followed by IF microscopy to visualize and measure analog-labeled tracts in replicated DNA. Sequential incorporation of the analogs enables identification and quantification of progressing replication forks, replication origins, and termination sites, depending on the labeling scheme. The measurement of individual DNA fiber length further provides direct information on the dynamics of replication fork progression. Because of these capabilities, DNA fiber spreading has established itself as an indispensable tool for studying DNA replication..

DNA fiber analyses have been hampered by the tedious and labor-intensive process of manually measuring the length of labeled fibers from IF images. While a handful of studies have attempted to automate this task using standard image analysis software pipelines, none of these solutions gained widespread adoption. This is likely because of their sensitivity to the quality and distribution of DNA fibers on IF images, leading to a high rate of errors, biases, and overall limited reliability. We previously proposed a DNA fiber measurement pipeline relying on classical computer vision techniques [11]. It involved preprocessing to identify sparsely populated regions, segmentation using a modified Canny edge detector, and diverse post-processing steps such as termination point reconnection and small blob removal. While conceptually sound, we observed that this approach tends to erroneously detect and segment too many fibers, especially in noisy or densely packed images. “DNA Stranding”, introduced by Li et al. [20], follows a similar approach and wraps the algorithm in a semi-automatic software providing a user-friendly interface. Users can refine segmentation results, offering a balance between automation and manual selection and measurement of fibers.

In the past decade, deep learning has established new standards in terms of segmentation performances [33]. While the use of these approaches for biomedical image analysis has gained considerable traction due to their improved generalization capabilities, no software solutions specifically adapted for DNA fiber analysis have yet been published in peer-reviewed journals.

Here, we present an experimentally validated, free, open-source, and user-friendly tool allowing the annotation and quantification of labeled DNA fibers from IF microscopy images obtained using the DNA spreading technique with two sequential pulses of nucleoside analogs. Our pipeline, called DNAi, integrates deep-learning based segmentation, various post-processing steps, and visualization in a unified framework. DNAi also proposes a length-based normalization strategy to calibrate automated outputs with manual expectations and a filtering module trained to detect false positives. The system is designed for usability and transparency, and provides a graphical interface that enables users to configure parameters and validate results interactively.

## Materials and Methods

### Cell culture

U2OS and HeLa cells (purchased from ATCC), were cultured in Dulbecco’s Modified Eagle Medium (DMEM; Gibco/Thermo Fisher) supplemented with 10% fetal bovine serum (FBS; Wisent), 5 mM L-Glutamine (Gibco/Thermo Fisher), and antibiotics (100 µg/mL penicillin and 100 µg/mL streptomycin; Gibco/Thermo Fisher). Cells were cultured at 37°C in 5% CO_2_. Cell lines were tested for the absence of mycoplasma contamination using the ZmTech Mycoplasma PCR detection kit (#M209001) and their identity was validated using STR Geneprint10 analyses (Promega, performed as a service from Genome Québec).

### siRNA Transfection

Cells were reverse-transfected with siRNA (Table 1) using Lipofectamine RNAiMax (Thermo Fisher) following the manufacturer’s instructions. The culture medium was changed after 24 hours, and DNA fiber assays were performed 48 hours after transfection.

**Table 1.**
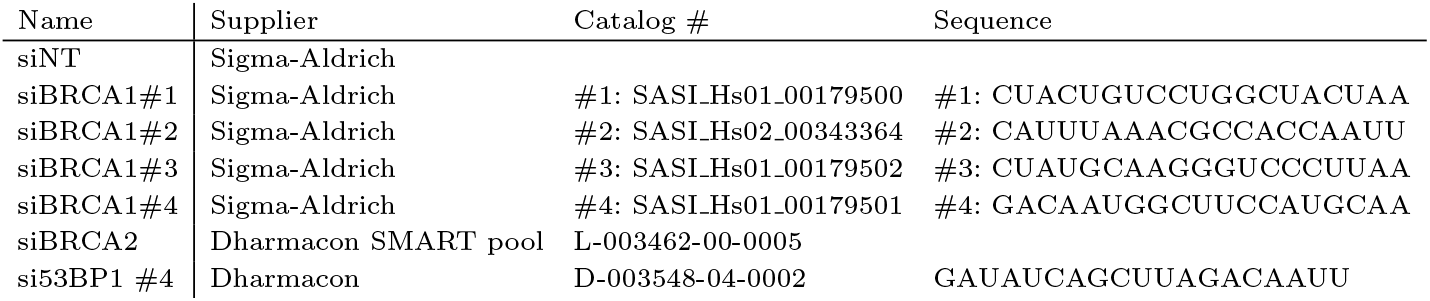
List of siRNA used in this study.

### DNA spreading assays

DNA fiber assays were performed as described previously with slight modifications [2]. Briefly, cells were seeded at a density 5 *×* 10^5^ cells per 6-cm dish and cultured overnight at 37°C in 5% CO_2_. Cells were pulse-labeled with 30 µM 5-chloro-2’-deoxyuridine CldU (Sigma-Aldrich) for 30 min, washed twice with PBS, and then incubated with 250 µM 5-iodo-2’-deoxyuridine IdU (Sigma-Aldrich) for 30 min at 37°C. Following labeling, cells were treated with 4 mM hydroxyurea (HU; Bio Basic Canada Inc.) for 4 hours. Cells were harvested by trypsinization, washed once with ice-cold PBS, and resuspended in PBS. Cells in suspension were spotted onto microscope slides, lysed with DNA lysis buffer, and tilted to stretch DNA fibers. Fibers were fixed in freshly prepared cold methanol/acetic acid (3:1, v/v) for 10 min, air-dried, and either stored at 4°C or processed immediately for IF. Slides were treated with 2.5 M HCl for 80 min, rinsed in H_2_O, and blocked with 5% BSA in PBS for 20 min at 37°C. Incorporated nucleotides were detected using rat anti-BrdU antibody recognizing CldU (Abcam, ab6326; 1:400) and mouse anti-BrdU antibody recognizing IdU (BD Biosciences, 347580; 1:25). Secondary antibodies were Alexa Fluor 594–conjugated goat anti-rat IgG (1:100; Thermo Fisher) and Alexa Fluor 488–conjugated goat anti-mouse IgG (1:100; Thermo Fisher). All antibody incubations were performed in a humid chamber at room temperature, followed by three washes in PBS-T (PBS with 0.05% Tween-20) and one wash with PBS. Slides were mounted with Immuno-Fluore mounting medium (MP Biomedicals) and stored at 4°C or –20°C in the dark until imaging.

### Microscopy platforms

Images were collected either using a 63×/1.4 oil objective lens on a widefield fluorescence Zeiss Axio Observer Z2 microscope with Zeiss Axiocam 820 monochrome sCMOS camera or a DeltaVision Elite System (GE Healthcare) using an Olympus 60×/1.42 as objective. Zeiss Zen blue software was used to capture Zeiss images and Resolve3D softWoRx-Acquire Version 6.5.2 for DeltaVision.

### Training data annotation

Training data were obtained by manually labeling ongoing and bidirectional replication forks (at a pixel level) on tiles extracted from whole-slide images. In total, we extracted 1,032 non-overlapping tiles of size 2048 *×* 2048 pixels. These were divided into a training set of 831 tiles and a testing set of 201 tiles, with 20% of the training set reserved as a validation set.

Four annotators participated in the annotation campaign, manually tracing all bicolor fibers they identified in each tile. To assess inter-annotator consistency, 20 tiles were independently labeled by all four experts. All annotations were performed using an in-house, open-source software called *LabelMed* (github.com/ClementPla/Annotator/). Annotators were instructed to trace fibers as accurately as possible using a pencil of fixed radius (3 pixels). In cases of ambiguity during color transitions (e.g., fibers appearing yellow), annotators were instructed to assign the pixel to the second label (i.e., green). An illustrative example of an annotated fiber is represented in Figure 1. In total, 6,232 fibers were labeled, with ratios ranging from 0.08 to 12.00 (median = 1.29).

**Fig. 1.**
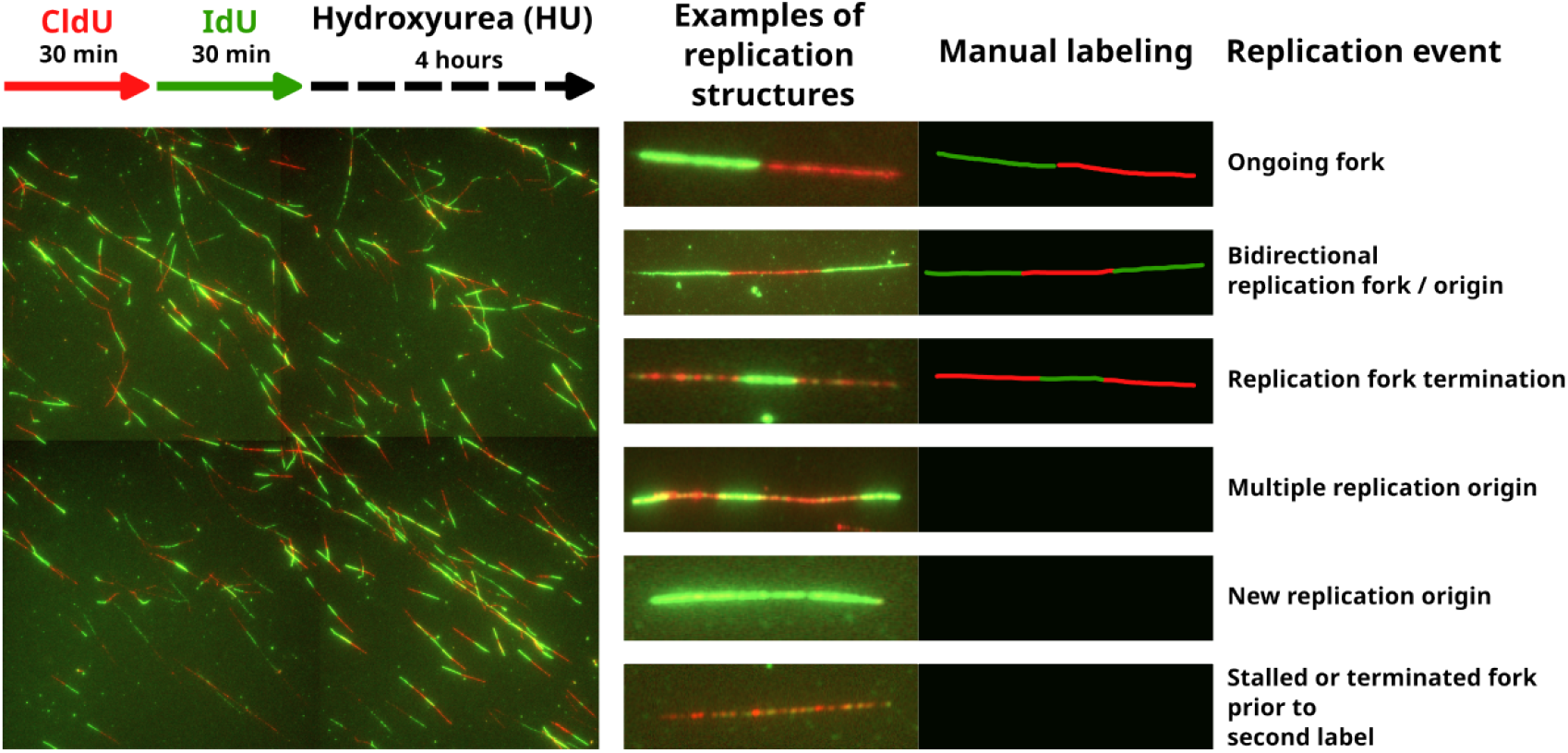
Example of manual fiber labeling. In cases of ambiguity in the analog signal (e.g., mixed colors or yellowish appearance), annotators were instructed to assign the segment to the second analog (IdU).

### DNAi pipeline

#### Preprocessing

To ensure uniformity and compatibility with our deep learning pipeline, several preprocessing steps were applied to all images. First, the spatial resolution was standardized by rescaling each image to a fixed scale of 0.26 micrometers per pixel. While this scale was fixed during training, the pixel size is configurable in the final tool to accommodate images from different acquisition systems. Next, histogram clipping was applied to suppress outlier intensities: values above the 99th percentile were clipped to reduce the influence of extreme brightness. Subsequently, all pixel intensities were normalized to harmonize brightness levels across the dataset. To further increase the robustness of the model and reduce overfitting, we applied a series of data augmentation techniques during training. These included random image rotations, horizontal and vertical flips, and rescaling, which collectively helped diversify the training examples without altering the underlying structures.

#### Segmentation model

To identify the most effective approach for DNA fiber segmentation, we evaluated several neural network architectures, ranging from well-established convolutional models to state-of-the-art transformer-based designs. Classical architectures such as the CNN-based U-Net [28] were included due to their proven performance in biomedical image segmentation tasks, with different encoders [31, 12, 29]. We also explored more recent architectures like SegFormer [32], which leverages a transformer-based encoder to better capture long-range dependencies and contextual information across the image. We used transfer learning, as each of the segmentation models had its encoder pretrained on ImageNet [7]. As a loss function, we used the Generalized Dice Loss, which we found to offer good performance, likely because of the very imbalanced class distributions. Each model was trained for 1000 epochs, with the AdamW optimizer, a learning rate automatically tuned before each training on 100 iterations, and a weight decay of 0.0001.

#### Post-processing

Following segmentation, post-processing steps were applied to refine the predicted fiber structures and enable accurate downstream analysis.

##### Junction Disentanglement

The first step involved skeletonization, where the segmented fibers were reduced to their 1-pixel-wide centerlines, preserving their topological structure while simplifying their geometry. This process is required to identify junctions (points where fibers intersect or branch) using a hit-and-miss transform. Specifically, each pixel neighborhood was tested against 12 predefined intersection templates. Once detected, each junction pixel and its 8 neighbors is temporarily masked out before reconnection. Any newly isolated component with a length of fewer than 10 pixels was deemed spurious and discarded (Figure 2-c). To reconnect the termination points, we traced the branch from the endpoint closest to the eliminated junction and gathered 15 pixels along the skeleton path to form a representative sample of the fiber’s trajectory at the crossing point. A least-squares linear regression was fitted to their coordinates (Figure 2-e). This provided a slope estimation, representing the directional angle of the fiber as it emerged from the junction. Based on both spatial and angular constraints, we then performed endpoint pairing to reconstruct the fibers. Two endpoints were considered a valid pair if their positions were within a maximum distance of 30 pixels and if the angle difference between their respective slopes was less than 20 degrees. When multiple candidate pairings were possible, the two with the shortest distance were selected. A simple heuristic was used for fiber coloring in the reconstruction process: if either of the connected junctions was labeled as second analog (green), a straight line was drawn with the label of the second analog; otherwise, the line was colored as first analog (red).

**Fig. 2.**
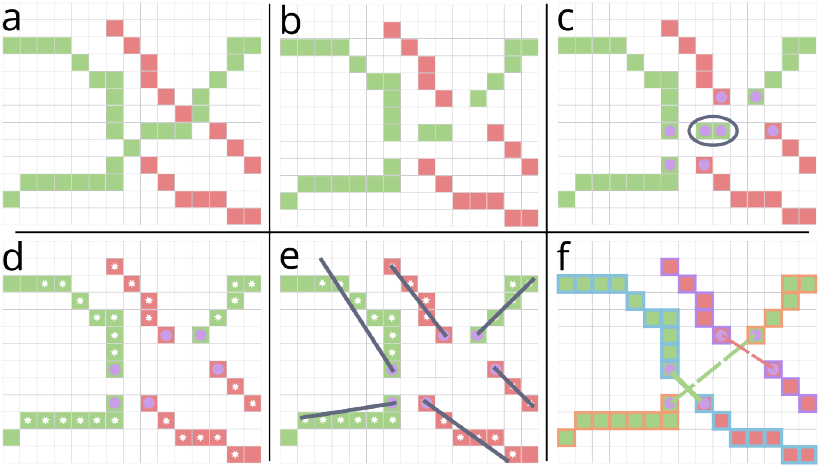
The post-processing steps taken to reconcile junctions in the segmentation: junctions identification and removal (a-b), island removal (c), endpoint orientation estimation (d-e) and fiber reconnection and clustering(f).

To eliminate noise introduced during segmentation, skeletonization and junction elimination, small connected components were then removed.

##### Error filtering

Although the segmentation models successfully detected a large number of fiber candidates, we implemented an additional optional filtering step to automatically remove aberrant segmentation towards improving the quality of the resulting quantifications with certain images. This solution relied on a secondary neural network that processed each candidate fiber individually. For every detected fiber, we extracted its bounding box and stacked the raw image patch with its corresponding segmentation mask as input channels. The classifier, trained as a binary decision model, distinguished between valid fibers and false positives. Given the relative simplicity of this binary classification task, the secondary network provided an efficient mechanism to automatically refine the segmentation output, ensuring that only high-quality fibers were passed to subsequent analysis stages. We trained the error detection model on 1,900 fiber thumbnails, of which 21.3% corresponded to manually identified errors. Following hyperparameter optimization, a ResNet50 architecture was selected as a suitable backbone for false-positive detection.

##### Final cleaning

In the final step, we discarded all fibers that are neither bicolor nor tricolor, as well as those located at the image borders to avoid including cropped fibers.

### Inference

To efficiently process large stitched mosaics of whole slides, we implemented a sliding window approach to enable full-image segmentation. In this strategy, each large mosaic is divided into smaller overlapping patches of 512 × 512 pixels, with a 25% overlap between adjacent patches. This overlap ensures continuity and consistency at the boundaries, reducing edge artifacts and improving the quality of the assembled segmentation. Following model inference on each patch, the individual predictions are reassembled into a full-resolution mask. To minimize the influence of border predictions, a Gaussian weighting function is applied during reconstruction. This approach assigns greater importance to predictions near the center of each patch, where the model’s output is typically more reliable, while softly blending overlapping regions to ensure smooth transitions across the image. Once this initial segmentation is obtained, the post-processing is done on the CPU. We use Numba to accelerate our code by precompiling Python into optimized machine code.

Finally, we implemented two more options within DNAi: ensembling and Test-Time-Augmentation (TTA) [24]. The former consists of several independently trained models whose predictions are combined, thereby reducing variance and improving robustness compared to a single model. TTA further enhances performance by applying a set of geometric transformations (horizontal flips, vertical flips, and 90° rotations) to each test image. Predictions are obtained for all augmented versions and then aggregated, which helps the model generalize better by averaging out orientation-dependent biases.

### Algorithmic efficiency

We report here the efficiency of different model families chosen among recent state-of-the-art, comparing the evolution of segmentation (Dice) and detection performance as a function of the number of floating-point operations (FLOPs) required for the forward pass of a 1024 *×* 1024 image. The detection metric is defined in line with the intuitive notion of precision: it is computed as the number of fibers detected both by DNAi and the human annotator (i.e., the intersection), normalized by the total number of fibers detected by DNAi. The size in terms of the number of parameters is represented by the respective radius of each point on the graph. We provide the weights of each model (Figure 3). All model variants are made available to the end-user, allowing a trade-off between computational cost and performance, adaptable to different hardware constraints.

**Fig. 3.**
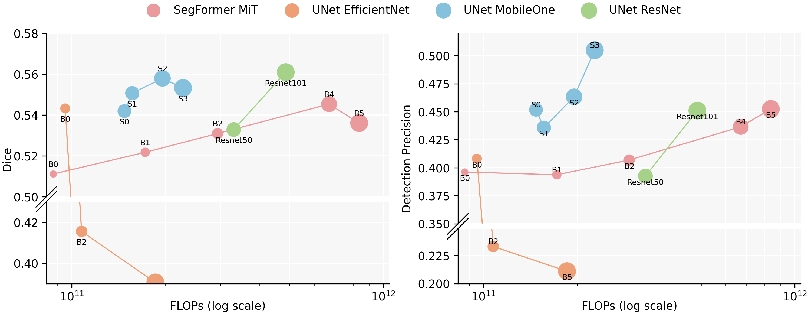
Comparison of several model variants, organized from smaller to larger architectures. The left panel shows segmentation performance, measured as the multi-class Dice score, plotted against the FLOPs (floating-point operations required to process an image of size 1024 × 1024). The right panel illustrates detection performance, defined as the number of fibers jointly identified by both the human annotator and DNAi, divided by the number of fibers detected by DNAi.

## Results

### Overview

We present a novel system called DNAi for the automated analysis of DNA fiber assays, leveraging deep learning for the segmentation of IF images. At the core of the pipeline is a neural network trained to identify pixels in IF microscopy images that correspond to DNA fibers containing consecutive, adjacent segments marked by nucleotide analogs(i.e., bicolor fibers). The network was trained using a collection of IF images generated from fork protection assays. Human cells (HeLa and U2OS) were exposed to CldU for 30 minutes, washed, then to IdU for 30 minutes, and finally incubated in hydroxyurea to induce replication fork stalling. This results in DNA fibers containing adjacent segments of CldU and IdU, representing progressing replication forks. In preparing our training dataset, we annotated only DNA fibers containing such adjacent CldU and IdU segments (ongoing and bidirectional forks). The total length (CldU+IdU) of annotated DNA fiber segments ranged from 2.34µm to 73.32µm, with individual segment length varying from 0.52µm to 49.4µm and ratio values from 0.08 to 12.0 with a median of 1.29. We used this spectrum of DNA fiber characteristics with the goal of generating a network capable of segmenting and measuring a wide variety of segment lengths and ratios, irrespective of the specific labeling conditions used by the experimenter. Nevertheless, the design of our training dataset implies that reliable detection, segmentation, and measurement of a DNA fiber requires it to contain two adjacent labels of different color in the final image.

We evaluated several neural network architectures, comparing their efficiency and performance for this task. As described in detail in material and methods, our pipeline includes post-segmentation tools for disentangling overlapping fibers and tracing individual ones to determine their length, identifying the fiber type, and IdU/CldU ratio. To further align automated outputs with human interpretation, we included an optional filtering module trained to detect false positives from the initial segmentation. The entire workflow is accessible through a user-friendly interface that supports configuration of key steps and interactive visualization of results.

Unless otherwise specified, all reported results are obtained using the ensemble models without applying test-time augmentation (TTA), followed by the error detection module to filter out unacceptable fibers. The impact of these two components is examined in more details in the following section.

### Inter-annotator variability and algorithm performance

DNA fiber IF images display highly variable quality and complexity due to factors related to both the acquisition itself (confocal or camera-based, sensor type, bit-depth, etc.) and the arrangement of DNA fibers (spatial density, alignment, overlap, etc.). On a set of 20 images independently annotated by four experts, we measured the number of fibers found by each and counted, for each annotator, the occurrence of fibers present in 1, 2, or 3 of the other annotations. The histogram in Figure 4 illustrates the inter-observer variability in the task of fiber detection. A fiber is considered common to two annotators if the intersection of the bounding boxes of the two annotations is at least 50% of their union. The data indicate that out of a total of 252 distinct fibers detected by human annotators, only 38 were detected by all four observers (i.e., 15.1%), and on average each annotator identified 21.1% of fibers that no other annotator found. This is consistent with our previously published results indicating that DNA fiber image interpretation typically leads to significant inter-annotator bias [11].

**Fig. 4.**
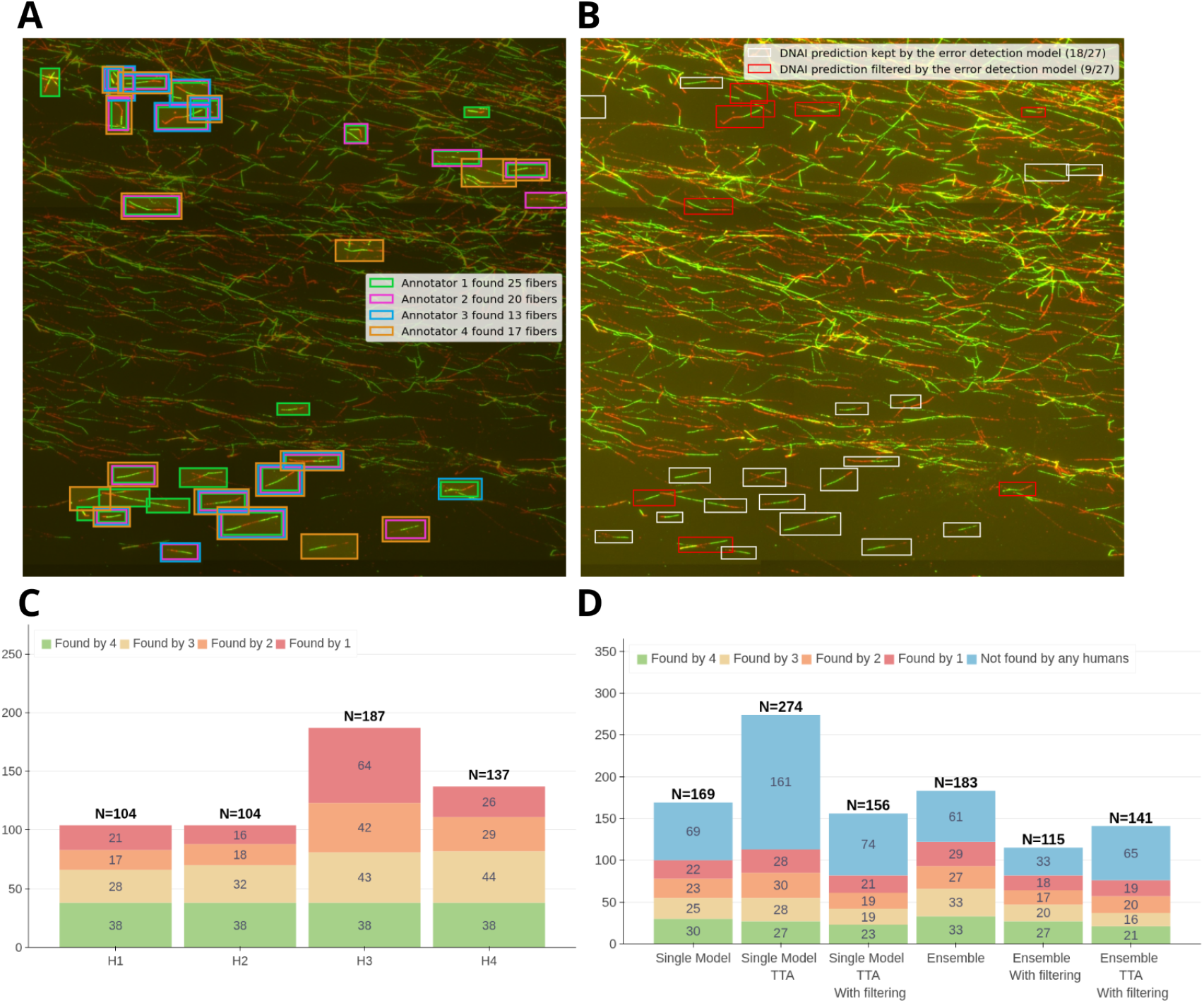
Inter-annotator variability. **A**: Example of a tile labeled by the four annotators. **B**: the prediction obtained with DNAi with or without the error detection model on the same tile. **C**: For each annotator *H*_*i*_, the plot shows the number of fibers that *H*_*i*_ found in common with zero, one, two or three of the others annotators. The count is obtained on the subset of the test set composed of 20 images labelled by the four annotators. **D**: Comparison of the fibers found by different variants of DNAi and the humans annotators, with or without the options we implemented.

The number of fibers detected by different variants of our models is presented in Figure 4. UNet-MobileOne S3 (single model in Figure 4-**D**), the model which we found had the highest precision on detecting fibers, segments a large number of fibers, of which a majority (59.0=2%) are detected by at least one human. This frequency decreases with TTA (41.2%), but in turns TTA vastly increases the number of fibers detected. In this configuration, the precision improves significantly by adding the error-detection model (52.6%). Ensembling models also improves the precision (66.7% alone and 71.3% with error detection), as represented in Figure 4-**D**.

To further validate the segmentation performance of the model, we used a test set consisting of a total of 201 images. This allowed us to quantify the ratio between the length of CldU and IdU labeled tracks in bicolor fibers.

Figure 4-**A**,**B** provides an example of the type of variability found in manual annotations and in the prediction made by the algorithm, with or without filtering.

Figure 5 shows the concordance of the ratios measured on the 809 fibers found in common (i.e., *IoU* ≧ 50%) between the human and DNAi. Among these fibers, we observe an excellent agreement between the human measurement and that of the algorithm, with a Pearson and Spearman coefficient equal to 0.88 (respectively 0.93), indicating that DNAi closely reproduces human segmentation. This supports the observation that, whenever humans and DNAi agreed on the detection of a fiber, their respective measurements (lengths and ratios) were similar.

**Fig. 5.**
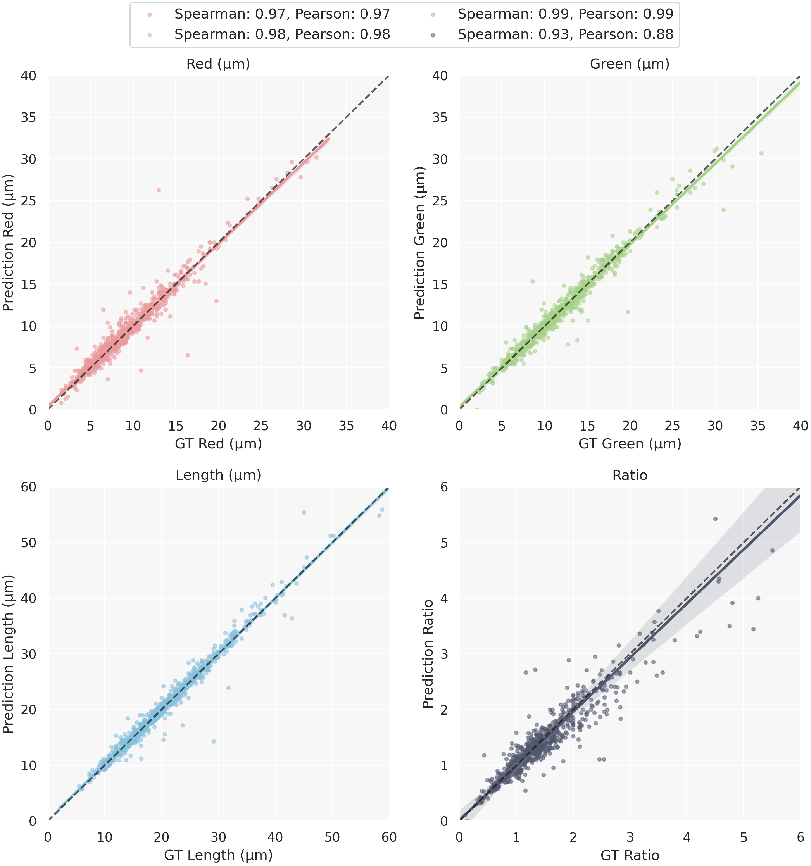
Comparison of fiber properties in the groundtruth (GT, the human annotator) and DNAi. The regression is computed only on fibers detected by both.

We next quantified how often such agreement occurs by evaluating precision and recall, comparing a same model with and without TTA and/or the error-filtering module. True Positives (TP) were defined as fibers in the predictions that also appeared in the ground truth, False Positives (FP) as predicted fibers absent from the ground truth, and False Negatives (FN) as ground-truth fibers missed by the predictions. The results (Figure 6) highlight the complementary trade-offs introduced by TTA and the error-filtering model. TTA increases recall but reduces precision, reflecting a higher number of fibers being detected, including some false positives. In contrast, the error-filtering model improves precision at the expense of recall, as fewer fibers are retained overall. The choice of configuration is situational. In many contexts, over-segmenting fibers is undesirable, making TTA less relevant. Moreover, TTA requires multiple inference passes, which significantly increases computational cost. However, in specific cases (for example when dealing with images outside the training distribution) DNAi may fail to detect a sufficient number of fibers. In such cases, TTA provides a simple yet effective way to boost detection rates.

**Fig. 6.**
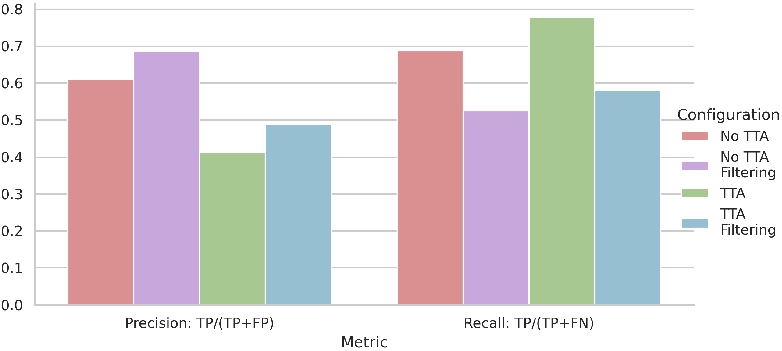
Precision and recall of DNAi on the 201 test images under different configurations. Results are shown for a single model combined with or without test-time augmentation (TTA) and/or the error-filtering module. TTA increases recall but reduces precision, reflecting a higher number of detected fibers, including some false positives. In contrast, the error-filtering module improves precision at the cost of recall, as fewer fibers are retained. These complementary trade-offs allow users to choose the configuration best aligned with their priorities.

### Assessing the performance of DNAi on datasets with limited annotations

The previous results were obtained under controlled conditions typical of machine learning protocols, using datasets annotated consistently for both training and evaluation. The limited size of the test set prompted us to extend the evaluation to more diverse scenarios. We validated DNAi on a larger set of images that were manually analyzed, and where only fiber-level ratio and label length measurements were available without any positional annotations on the images themselves. Images were processed and DNA fiber were manually measured using Image J (Fiji). We collected data from 27 biological condition, representing a total of 400 stitched images, conducted under various genetic silencing leading to, according to manual analysis, median IdU/CldU ratios between 0.82 and 1.71, min/max ratio values of 0.18 and 5.68 and overall length ranging from 2.93µm to 104.04µm.

Each experiment included manual measurements of approximately 200 fibers, resulting in a total of 5,881 manually measured fibers. In contrast, DNAi processing of the same dataset identified 32,864 fibers, as no upper limit was imposed on the number of fibers analyzed per experiment. The detected fibers spanned a wide range of lengths, from 1.17 µm to 102.96 µm. To ensure that comparisons were not biased by this discrepancy in sample size, we also implemented a simple calibration strategy: instead of considering all detected fibers, we selected a fixed subset (e.g., 200) whose lengths were closest to the average fiber length in the predicted population. This choice is motivated by the hypothesis that outliers in ratio measurements are more likely to arise from very short or long fibers, which human annotators would typically disregard. The approach is unsupervised, facilitates a fair comparison, and preserves the integrity of automated analysis. Overall, we observe excellent agreement, both in terms of linear correlation and relative ranking across conditions. Nevertheless, DNAi shows a systematic bias toward higher ratio estimates. As shown in Figure-7-**C**, the calibration strategy reduces the systematic overestimation and brings DNAi measurements closer to human values. In the DNAi user interface, an option is available to enable this calibration.

Figures 7-**B**,**C** show the comparison before and after the calibration step with a regression analyses comparing the median ratio values from manual and automated measurements. Without filtering (Figure 7-**B**), DNAi exhibits a systematic bias toward higher ratios. Upon inspection, we believe the disagreement arises from the interpretation of the yellow part of the fibers, indicating a transition between nucleotide analogs, particularly in the case of short fibers. After calibration (Figure 7-**C**), this bias is substantially mitigated, resulting in stronger correlation and improved consistency with manual assessments. These results underscore the importance of post-processing strategies in bridging the gap between automated and expert-driven analyses.

**Fig. 7.**
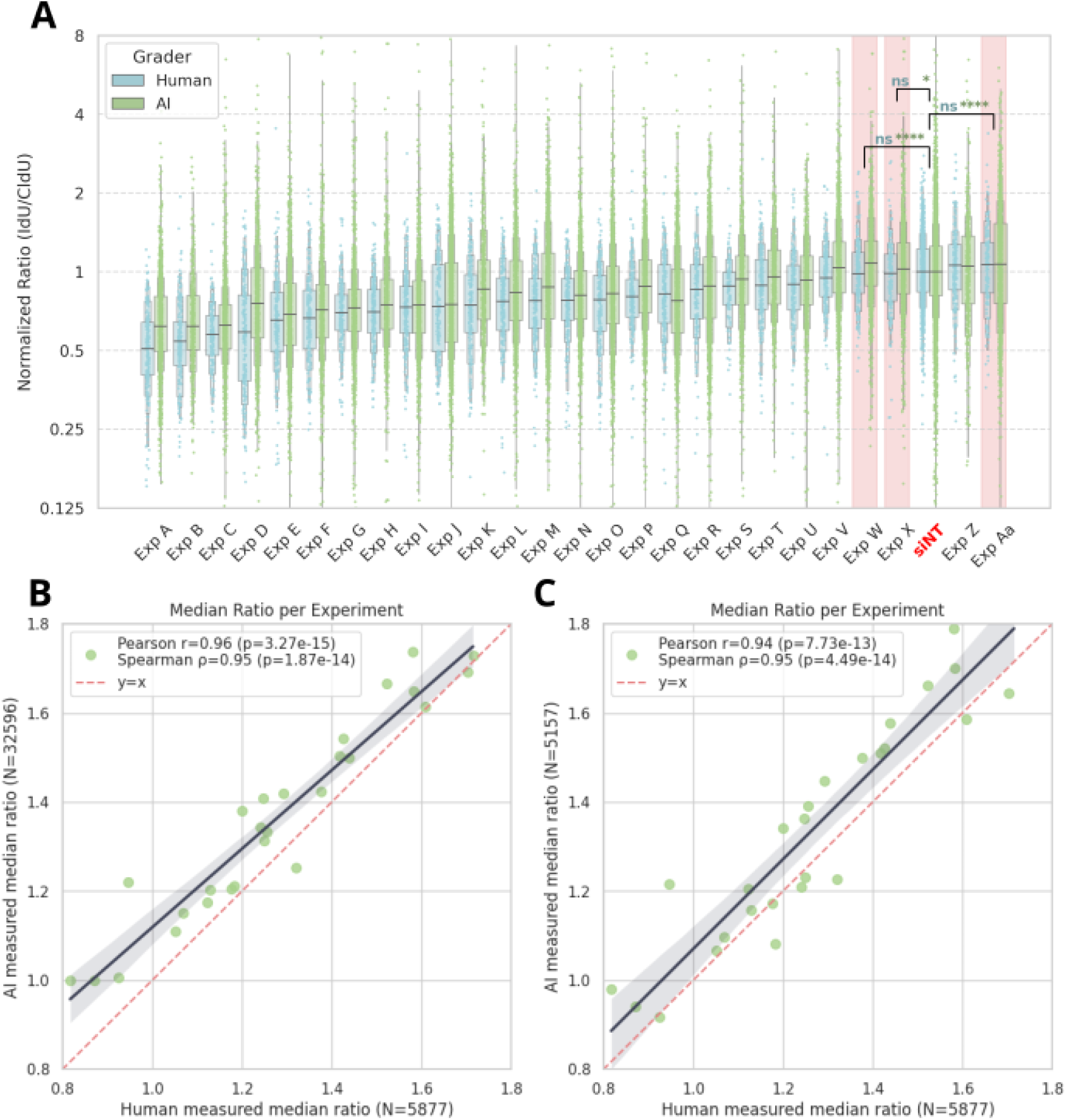
We evaluated DNAi across 27 biological conditions, corresponding to a total of 400 stitched images. For each condition, a human annotator reported the ratio of over 200 fibers, providing a robust reference for comparison. (**A**) Normalized ratios are ranked according to the human median. The siNT condition is used as a control. In pink, conditions where DNAi detected a statistically significant difference relative to the control whereas the human annotator did not. For the remaining 23 conditions, DNAi and the human annotator showed full agreement regarding the presence or absence of a significant difference compared to siNT. (**B**) Regression of the median ratios measured by DNAi against those reported by the human annotator. (**C**) Calibration procedure applied to reduce the systematic bias toward higher ratio estimates.

### Robustness to image degradation

To assess the robustness of the DNAi ensemble under suboptimal imaging conditions, we conducted a series of *in silico* experiments simulating two prevalent forms of image degradation. First, we applied Gaussian blurring with increasing standard deviation *σ* to emulate out-of-focus acquisition. Second, we introduced additive Gaussian noise with progressively higher variance to simulate random pixel-level disturbances.

Model performance was evaluated by monitoring changes in fiber-level ratio measurements, providing insight into the ensemble’s tolerance to real-world imaging. Figure 8-**A** summarizes the results on a representative high-resolution image containing approximately 300 fibers (obtained from DNAi, without perturbation). We compared three configurations: a single model, an ensemble, and an ensemble augmented with TTA.

**Fig. 8.**
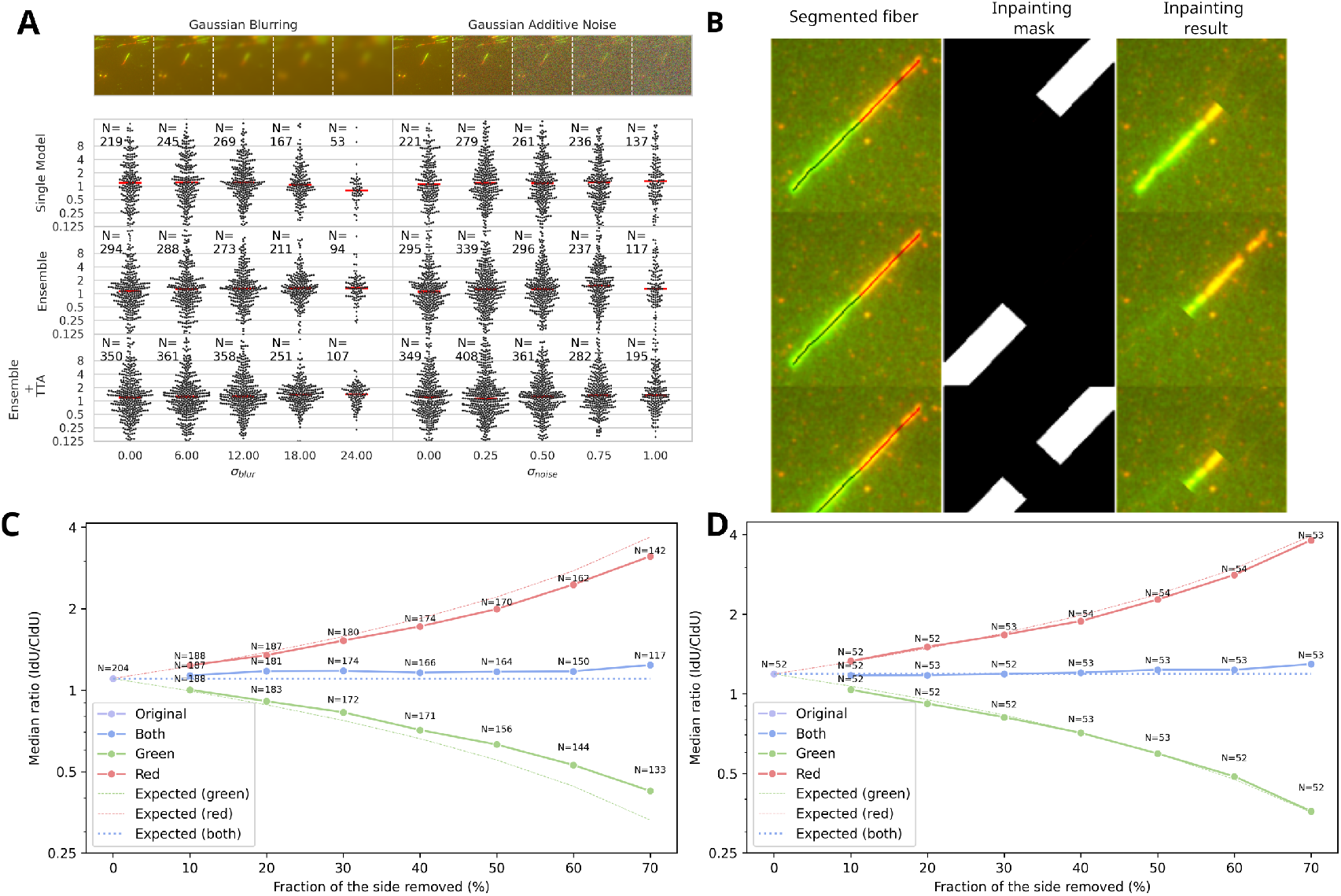
(**A**) Robustness of DNAi to image degradation. Images were progressively degraded using Gaussian blur (left) or additive noise (right). For each level, we report the median distribution of fibers detected by a single model, an ensemble, and an ensemble with TTA. All models remain consistent across degradations, though fewer fibers are detected as quality decreases. TTA improves sensitivity, enabling DNAi to recover more fibers under challenging conditions. (**B–D**) Fibers were artificially shortened from one or both sides by a predefined fraction, with the cut regions reconstructed by inpainting. (**C**) shows that DNAi’s measured ratios follow the expected trajectory, though slight mismatches occur because some fibers become too small to be consistently detected. (**D**) addresses this by computing the median only over fibers present at all cut levels, restoring close agreement with the expected trajectory.

While all configurations exhibit stability under mild perturbations, we observe a marked decline in the number of fibers detected under severe degradation. However, notably, the ensemble configuration maintains consistent ratio measurements despite the reduced fiber count, indicating resilience in preserving biologically relevant metrics even when detection sensitivity is compromised. It is important to note that, in these experiments, the error detection model was deliberately deactivated. This decision stems from its tendency to classify as false positives any fibers detected in regions affected by high levels of degradation. Such behavior, while intended to improve precision, can lead to excessive rejection of valid fibers under challenging imaging conditions. Indeed, we opted to keep this setting optional as users may wish to improve detection sensitivity when image quality is reduced.

Quantitatively, from the original images to the most blurred ones (*σ* = 24), we observe variations in the measured median ratio of –32.6%, +18.1%, and +18.4% for the single model, the ensemble, and the ensemble with TTA, respectively. Under noise degradation, the variations are more modest, at +10.5%, +13.9%, and +8.1%. These values should be interpreted together with detection performance: under blurring, the number of detected fibers drops substantially—by 75.8%, 68.0%, and 69.4% for the three configurations. In contrast, additive noise has a milder effect, with fiber detection decreasing by 37.0%, 60.3%, and 44.1%. While these drops in detection may appear substantial, they underscore the robustness of all configurations in preserving the main measurement of interest, the fiber ratio. Going a step further, this suggests that the fibers for which the network is consistently confident (those still segmented even under strong degradation) are representative of the overall fiber population.

### Simulation framework for algorithmic bias and ratio Control

Results confirm the overall consistency of DNAi predictions with human measurements. However, when discrepancies arise, it is unclear how to determine which source (the human observer or the algorithm) is biased. Thus, we aimed to evaluate the algorithm’s performance on fibers with a wide range of nucleotide analog ratios, in order to highlight its ability to generalize across a controlled spectrum.

To this end, we designed an *in silico* experiment that allows precise control over the ratio of fibers present in a slide, by artificially modifying fiber ratios in real images. Starting from an initial segmentation of the fibers, we shortened the skeleton of the segmented mask for the first analog, the second, or both. To propagate this modification onto the image, we generated a mask that encompasses the targeted cut region. Within this masked region, we applied a simple inpainting technique: each pixel was sampled from the area surrounding the fiber. Our method requires only two input parameters: the targeted analog (red, green, or both), and the fraction of its length to be removed. Figure 8-**B** illustrates the outcome of this technique on the same fiber for different parameter values.

Given an initial ratio *α* and cut fractions per analog *C*_*r*_ and *C*_*g*_ (for red and green respectively), the measured ratio evolves as:

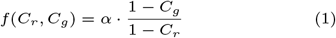

The outcomes of the in silico experiments (figure 8-**C**,**D**) closely follow the theoretical expectations. The measured ratios evolve in line with the predicted values across all tested configurations. This strong agreement demonstrates that the model reliably segments fibers regardless of their individual lengths or ratios. Importantly, even when fibers are systematically shortened from one or both sides, the algorithm consistently recovers the correct population-level ratio. These results confirm the model’s capacity to generalize and maintain accuracy across a broad spectrum of fiber compositions.

### Detection of biologically relevant differences in IdU/CldU ratio

One mechanism by which cells tolerate blockage of DNA polymerase activity is fork reversal [1]. During this process, the newly synthesized DNA strands anneal together, forming a four-way structure commonly referred to as a “chicken foot.” This exposes a double-stranded DNA end that is susceptible to nucleolytic degradation. So-called “Fork protection” factors, including BRCA1, BRCA2 and 53BP1, safeguard against this degradation thereby preserving nascent DNA integrity [21].

To evaluate the biological relevance and sensitivity of DNAi in detecting replication stress phenotypes such as nascent DNA degradation caused by loss of BRCA1, BRCA2, or 53BP1, we compared fiber-level ratio distributions between control conditions (siNT) and gene-silenced conditions targeting either BRCA1, BRCA2 or 53BP1 using siRNA. Cells were exposed to nucleotide analogs as described before (30 minutes CldU, wash, 30 minutes IdU), followed by a 4h treatment with hydroxyurea to block replication fork progression. Nucleolytic degradation of the nascent DNA containing the second analog during hydroxyurea-induced fork stalling in cells lacking BRCA1/2 or 53BP1 is expected to cause a reduction in the IdU/CldU ratio compared to control cells. We compared manually measured fibers ratios with those obtained using DNAi, using the length-based calibration strategy described in section 4.3.

Figures 9-**A** and 9-**B** illustrate the distribution of ratio measurements for each condition. As expected, we observed a statistically significant shift in the distribution under BRCA1, BRCA2, or 53BP1 silencing, consistent with known effects on nascent DNA stability at stalled forks, in both the manual and DNAi measurements. These findings demonstrate that DNAi not only replicates the trends observed in expert annotations but also reliably captures the effects of gene knockdowns on fiber ratios. The consistency of statistical significance across both annotation modalities underscores the robustness of the model and its ability to yield biologically meaningful insights.

**Fig. 9.**
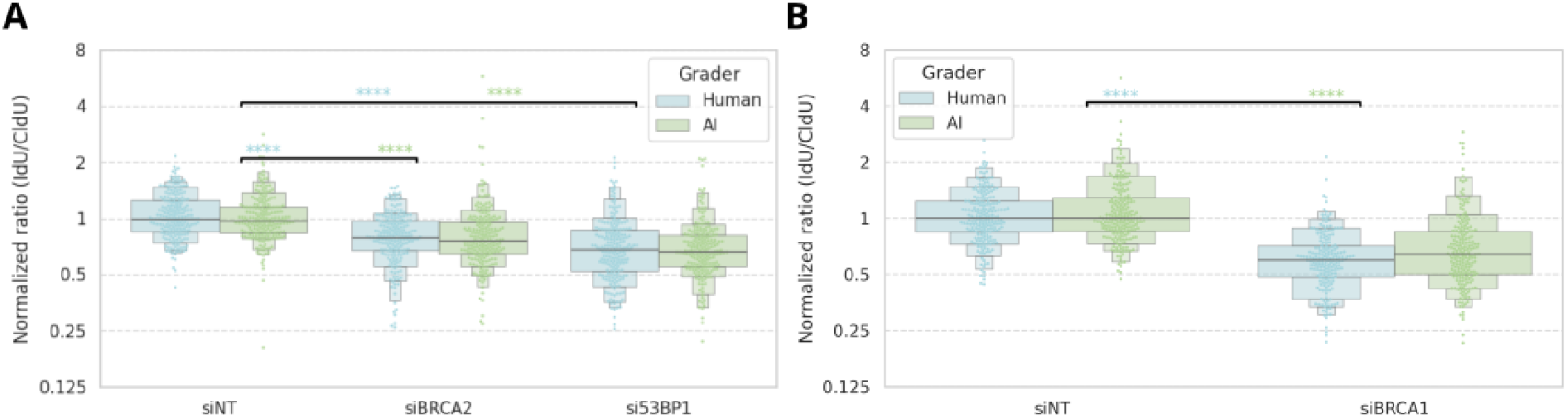
Distribution of fiber ratio measurements under different experimental conditions. (A) Comparison of IdU/CldU ratio measurements between siNT, siBRCA2 and si53BP1. (B) Comparison of siNT with siBRCA1. In both panels, distributions are shown for fibers annotated manually by human experts and detected automatically by DNAi. Statistical significance was assessed using a Mann–Whitney U test, confirming a highly significant difference between the control and each experimental condition (*p <* 0.001) for both manual and DNAi measurements.

### Generalization to other acquisition device

Images used for model training and for the tests described in the preceding figures were acquired on a Zeiss Z2 microscope (see material and methods). We evaluated its performance on images acquired with a microscope from a different manufacturer (DeltaVision, which is based on an Olympus platform; see section 3.4) over various experiments (Figure 10). Visually, contrast, pixel size and overall signal-to-noise appears to differ between images acquired using DeltaVision or Zeiss systems, which make the following experiments a good measurement of the robustness of the model to out-of-domain inference. The data indicate that DNAi is able to reliably captures relative trends in fiber ratios across experiments, even if minor shifts occur compared to the human counting that we did not observe at this scale on Zeiss images.

**Fig. 10.**
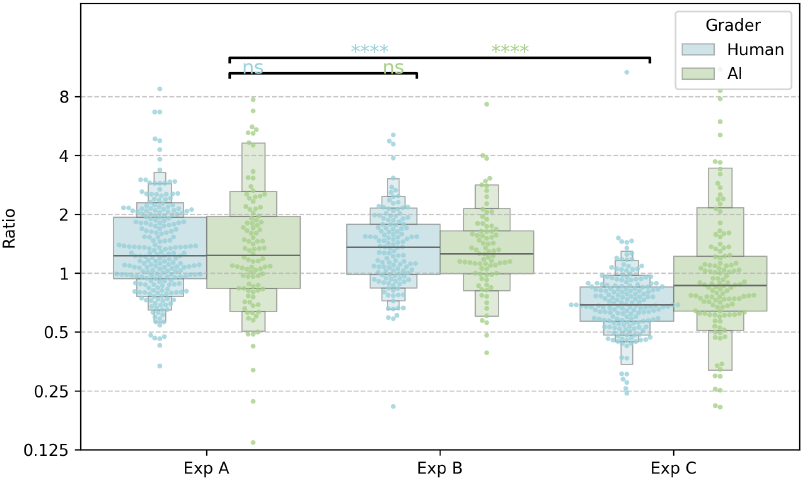
DeltaVision experiment. DNAi reproduces the statistically significant differences observed by human annotators across multiple experimental conditions. While the direction and significance of the differences are fully consistent, the absolute median fiber ratios show slightly larger deviations from human measurements compared to the results obtained on the Zeiss microscope. This indicates that overall DNAi reliably captures relative trends in fiber ratios across experiments, even if minor shifts occur depending on the imaging platform.

### Graphical User Interface

To ensure usability, the entire pipeline is integrated into a graphical user interface (GUI) aimed at non-technical users. The interface allows configuration of all analytical parameters and supports both single-image and batch processing. Visualization tools include an interactive display of original and segmented images, fiber bounding boxes, and a tabular view of individual fiber metrics. Users can manually exclude fibers from analysis and generate population-level statistics using built-in calibration and regression features. To ensure cross-platform compatibility, the UI was developed as a web application running locally, allowing its use in any modern web browser.

In summary, the software is composed of the following configurable modules:

- Flexible image loading and preprocessing, including decoding of.cvi,.dv,.tiff,.jpeg, or.png formats. Preprocessing is configurable, allowing specification of pixel size, marker ordering and bit depth. Users can load dual-channel images directly or load each channel separately.
- A segmentation module offering a several pre-trained architectures with options for ensemble inference and/or TTA.
- An error-detection module that can filter false positives from the segmentation model with adjustable aggressiveness.

The viewer supports multiple visualization modes (bounding boxes, plain strokes, or animated strokes) and allows users to modify the status of a fiber (valid or invalid) with a simple click. Image-level statistics can be exported as a.csv file directly from the table view, which also enables detailed inspection of individual fibers. The GUI also allows users to adjust the prediction threshold applied to convert the probabilistic output of the segmentation network into discrete predictions. Since the model is a three-class architecture (background, first analog, second analog), the default threshold is inherently set at 1/3. Lowering this value enables the model to extract more fibers from the probability map, at the potential cost of increased false positives. In addition, the GUI includes a batch analysis tab that sequentially processes all loaded files. The results can be exported as a.csv file or displayed as a swarm plot, with filenames automatically used to group images by experimental condition.

## Discussion

DNAi is a suite of software tools that can be used either via scripts or a graphical interface. It is intended as open-source software that is easy to use and significantly improves the productivity of biologists performing DNA fiber assays. Our study demonstrates the technical feasibility of designing an AI for automatic segmentation of DNA fibers. Importantly, advances in microscopy enable the generation of increasingly large datasets from DNA fibers assays. DNAi and similar computer vision software are expected to facilitate efficient analysis of these datasets, replacing the manual counting of a handful of fibers with more accurate and reproducible analyses.

Our software not only expedites the quantification of DNA spreading experiments but also leads, in general, to a vastly increased number of detected DNA fibers per image. This is expected to improve the interpretation of experimental results: indeed, in our experience DNA spreading can lack uniformity in quality of labeled segments from one microscopy field to another. This can lead to biased results when experimenters only count a limited number of fibers (usually around n=200) from a few microscopy fields. In addition, our previous data [11] and those presented here (Figure 4) clearly demonstrate that the selection of fibers to be measured varies significantly between human observers, further biasing image interpretation. While our deep learning model is likely to present biases since it was trained with manually segmented images, it uniformizes such biases across experimental conditions. We expect that this will improve the overall reliability in the quantification of relative differences in DNA fiber length and nucleotide analog ratios.

### Limitations

Despite its overall robustness, DNAi has several limitations that must be acknowledged. As mentioned before, the models were trained on a relatively constrained dataset, primarily composed of images resulting from one type of labeling scheme (two nucleotide analog pulses of equal duration followed by hydroxyurea treatment), which were acquired from a single microscope and annotated by a limited number of experts. This introduces potential biases and limits generalizability. To mitigate this, we incorporated variability in acquisition conditions, applied data augmentation, and used ensemble inference strategies. Moreover, we selected datasets which presented an approximately equal number of experiments whose median IdU/CldU ratios were either smaller, equal, or larger than 1, and containing fibers with ratios and length values significantly deviating from the median. In our hands, DNAi is able to detect a variety of DNA fiber length and analog ratios reflecting typical DNA spreading labeling schemes and results, and does not appear to be specifically biased toward any type of ratio or DNA fiber length. Nevertheless, users should remain vigilant and verify results when using DNAi with images originating from labeling schemes or experimental protocols that significantly deviate from those used to train our model.

Our model was trained and validated using the DNA spreading technique with a two nucleotide analog labeling scheme, i.e., resulting in two-color IF images. As a result, the software does not currently detect single color DNA fibers. Our validation experiments also indicate that fibers containing more than two segments, e.g., those deriving from converging forks or replication origins, are not detected and segmented with as much efficacy as those containing only two segments. This is expected to reflect the inherent paucity of such fibers in the training dataset. Because of this, other deep learning models would likely need to be trained to robustly segment and quantify DNA fibers containing more than two labeled segments. We also note that while DNA combing and DNA spreading are conceptually related, the resulting microscopy images reveal that DNA fibers observed using these methods differ noticeably with regards to fiber length, shape, and continuity. While we have not tried to optimize DNAi for DNA combing applications, dedicated deep learning models will probably need to be trained specifically for this application.

While our data and validations indicate that DNAi is robust to a wide variety of imaging conditions, visual examination reveals that segmentation errors persist in the DNAi output, particularly when using images that are of lower quality or displaying densely packed fibers. While these errors affect individual measurements, analyses presented in the current work show that at the population level, their limited frequency limits their impact on distribution metrics. To address over-detection, which we observed across several models, we added an optional secondary error-detection module that considers both segmentation and local image context. This module improves alignment with human annotations in many cases but can reject fibers from imaging sources outside the training distribution. For example, we have observed empirically that images obtained using a DeltaVision microscope tend to result in a higher rejection rate than those generated using a Zeiss Z2 system. For this reason, we provided the option to disable this step via the graphical interface, which can improve segmentation with certain sets of images. This filtering model, while useful for removing obvious false positives, suffers from the same interpretability limitations as most discriminative neural networks. It may reject valid fibers without obvious justification, especially under conditions of image degradation. In future work, we aim to extend the pipeline with an image quality assessment module to guide users and improve reliability across diverse acquisition conditions.

These observations lead us to advise using the software through its graphical interface, which allows users to visually confirm that the model’s output aligns with human expectations. The GUI also exposes a range of adjustable parameters (e.g., pixel size, marker indexing order, model selection, detection sensitivity), enabling users to tailor the pipeline to their specific experimental setup. Visualization modules at the image, fiber, and distribution levels further support manual validation and correction.

In summary, our work demonstrates the feasibility of training deep learning models for DNA fiber quantification tasks. We anticipate that this and future software solutions will improve throughput and diminish inter-user variability and biases in the interpretation of such experiments, thereby improving the robustness and reliability of the results.

## Data availability

The data underlying this article (images and the corresponding masks used to train the models) are available at the following link: https://doi.org/10.5281/zenodo.17237308[26]. The complete code for training and using DNAi is available in the following github repository: ClementPla/DNAi. Models weights are automatically downloaded on usage from a repository on HuggingFace.

## Author contributions statement

C.P. contributed to conceptualization, data curation, formal analysis, investigation, methodology, validation, project administration, software, visualization and writing.

Y.M. contributed to conceptualization, data curation, formal analysis, investigation, methodology, validation, visualization, and writing.

R.D. and M.C. B. contributed to funding acquisition and supervision.

S.C. contributed to conceptualization, supervision, project administration and writing.

H.W. contributed to conceptualization, obtained funding, wrote and reviewed the manuscript.

## Acknowledgments

This work is supported by funds from the Natural Sciences and Engineering Research Council of Canada to H.W. (RGPIN-2019-05082 and RGPIN-2025-04612) and S.C. (RGPIN-2021-03330) and from the Line Chevrette foundation to C.P. S.C. acknowledges support from the WOCO foundation.

